# Theory of antiparallel microtubule overlap stabilization by motors and diffusible crosslinkers

**DOI:** 10.1101/692962

**Authors:** Manuel Lera-Ramirez, François J. Nédélec

## Abstract

Antiparallel microtubule bundles are essential structural elements of many cytoskeletal structures, for instance the mitotic spindle. In such bundles, neighbouring microtubules are bonded by specialised crosslinkers of the Ase1/PRC1/MAP65 family that can diffuse longitudinally along microtubules. Similarly, some kinesin motors implicated in bundle formation have a diffusible tail allowing them to slide passively along microtubules. We develop here a theory of two microtubules connected by motors and diffusible connectors, in different configurations that can be realized experimentally. In all cases, the microtubule sliding speed derived analytically is validated by stochastic simulations and used to discuss recent experimental results, such as force generation by kinesin-14, and overlap stabilization by Ase1. Some systems can produce steady overlaps that are determined by the density of crosslinkers on the microtubule lattice. This property naturally leads to robust coordination between sliding and growth in dynamic bundles of microtubules, an essential property in mitosis.

## 1 INTRODUCTION

Arrays of parallel microtubules are indispensable in cells, appearing for instance in mitotic spindles [24], neuronal dendrites [28], or marginal bands of blood platelets [11]. Observed in cross-section by electron microscopy, these arrays have a regular organisation that is composed either of square, triangular [24], or even hexagonal unit cells [21]. These bundles are formed by specialised molecular crosslinkers that mechanically connect adjacent microtubules. Some crosslinkers only bind when the microtubules are oriented in the same direction [17], while others bind only when the microtubules are antiparallel [24]. Molecular motors can also connect microtubules and slide them relative to each other, resulting in overlap shortening, and these changes can be opposed by crosslinkers. Overlaps formed *in vitro* with stabilised microtubules can reach an equilibrium length, that arises from the interplay between molecular crosslinkers and motors. Bundles found *in vivo* often include dynamic microtubules, but their overlaps can nevertheless reach a steady state length. In the case of central spindle overlap, for instance, microtubule plus ends elongation leads to additional sliding, whereby the overlap length appears to remain constant, preserving the mechanical connection between the microtubules [16]. The resulting slow separation of spindle poles is thought to be necessary to complete mitosis [18]. Unravelling the mechanisms by which microtubule overlaps are stabilised and regulated is essential to understanding mitosis and other key processes in cell biology.

In recent years, a number of systems involving diffusible crosslinkers and motors have been studied *in vitro*. Importantly, this work showed that two microtubules were sufficient to form a stable antiparallel overlap, if the correct kind of motor was used. Particularly, kinesin-14 was shown to lead to stable overlaps [1], in contrast to kinesin-5, which moves antiparallel microtubules apart at constant speed [19]. Noteworthy, the kinesin-14 family members Ncd and HSET, in addition to a motor head, contain a diffusible tail [2], making them able to form asymmetric connections with motor and diffusible head bound to different microtubules. Interestingly, Kinesin-14 can only reach sub-picoNewton forces when crosslinking antiparallel microtubules [10], even though its motor domain is able to exert picoNewton forces *in vitro*. This suggests that the diffusible tail of kinesin-14 limits the force exerted by its motor domain.

The crosslinkers of the MAP65/Ase1/PRC1 family preferentially crosslink antiparallel microtubules [6], and can diffuse longitudinally along single microtubules and microtubule overlaps [7]. Fission yeast cells expressing excess Ase1 exhibit slower mitotic spindle elongation [16], [8]. In HeLa cells, PRC1 is required for the stabilisation of the anaphase midzone [29], and has recently been observed to locate to the bridging fibres connecting sister k-fibres, suggesting that this protein may also have a role during metaphase [15]. *In vitro*, diffusible crosslinkers can oppose sliding by molecular motors [1], [9], [26]. Being passive in nature, one might have expected the force required to move a head to be proportional to the sliding speed. It was found however that the resistance to sliding can increase dramatically with the density of the molecules on the antiparallel overlap [1], suggesting the existence of a critical density of crosslinkers above which the system jams. These crosslinkers are also able to widen an overlap *in vitro*, in the absence of any motor [9], leading to the idea that they could be regarded as a gas confined within the overlap. Such phenomena were understood by considering the discrete nature of the microtubule lattice with its well known periodicity of 8 nm, given that only one head, at most, may bind to a tubulin heterodimer. In these experiments, diffusible crosslinkers remain associated preferentially with the overlap region, where the two binding domains can be bound. Statistically, overlap extension creates more possibilities for the crosslinkers to bind, resulting in an entropic pressure effectively pushing the microtubules ends apart. Inversely, when sliding results in the densification of the crosslinkers in the overlap, the resistance to sliding increases, eventually reaching a steady state overlap length, which can remain stable for several minutes [1], [9], [27], [4].

Entropic expansion was previously modelled using both an analytical model and a computational model (lattice-based stochastic simulations) [9]. Interestingly, while the analytical model did not quantitatively match the experimentally observed behaviours, the computational model did, as it exhibited a drag that increased exponentially with the number of crosslinkers. Discrete models are arguably more complicated to analyse, as they rely on many assumptions, any of which could greatly influence the dynamics of the system. We extend their analytical theory, showing good agreement with their experiments. Diffusible crosslinkers were modelled so far without active force generators, following in vitro conditions. However, in the cellular context, they most likely operate together with molecular motors. Whether some coordination mechanism is needed to form stable overlaps, or not, has not been yet been theoretically examined.

Aiming to understand how motors and diffusible microtubule binders can be combined to form stable overlaps, we study here four related systems (Fig. 1) inspired by the *in vitro* work pre-cited. All systems contain two antiparallel microtubules and we only consider their motion in one dimension. We also presume a constant overlap length, assuming that microtubule growth matches the sliding exactly. This point will be discussed. The four systems differ in the way the diffusible and motor heads are associated to form connecting molecules. In system A, crosslinkers bind and unbind but do not diffuse along microtubules, while sliding is produced by bivalent motors. System B is similar, except that the crosslinkers can diffuse along microtubules and never unbind. It was realised *in vitro* using PRC1 and kinesin-5 [20]. System C corresponds to experiments using Kinesin-14 [10], in which the sliding is produced by motors composed of a diffusible tail and a motor head, without crosslinkers. Finally, in system D, diffusible motors pull against diffusible crosslinkers. This was explored with Kinesin-14 and Ase1 [1], [9]. Another system in which Kif4A motors directly pulled on PRC1 crosslinkers was modelled previously by us [4], and thus omitted here. These systems offer gradual complexity and different outcome. After defining a common set of assumptions, we predict the sliding speed of the microtubules in each system.

**Fig 1:**
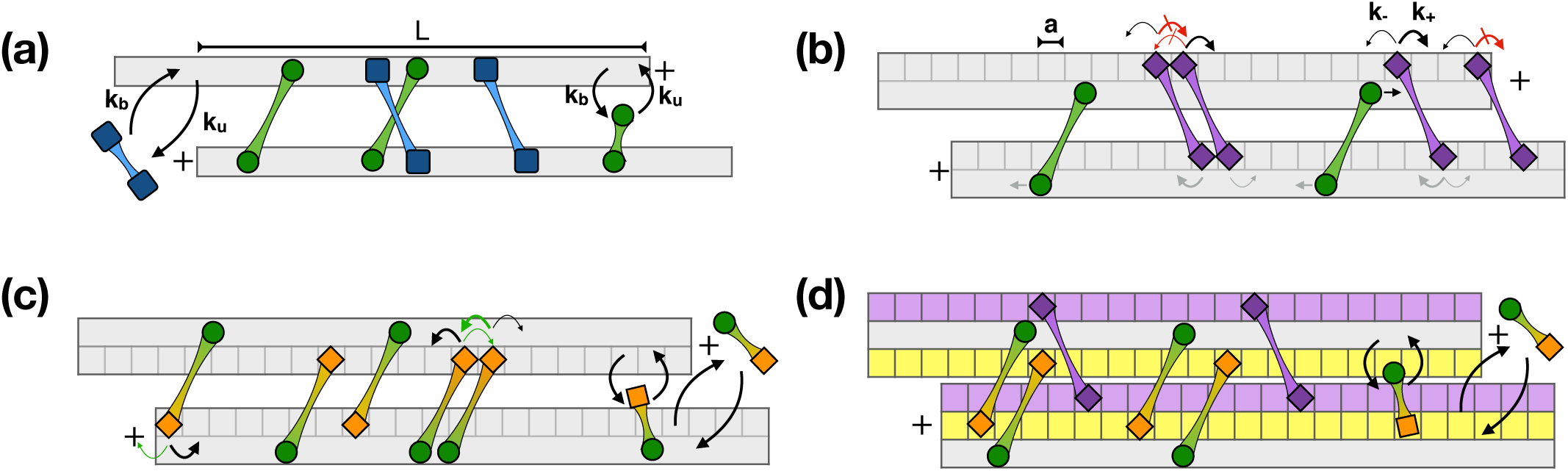
Schematics of the modelled systems. Systems containing two antiparallel microtubules, arranged with an overlap length *L* that is constant because microtubules grow at the required speed to compensate exactly for the sliding. **(a)** In system A, microtubule sliding is determined by bivalent motors (green circular heads) and crosslinkers (blue square heads). These crosslinkers bind and unbind but do not slide along microtubules. Motor heads move actively towards the plus ends and tend to reduce the overlap. Crosslinkers resist this motion until they unbind. **(b)** System B combines bivalent motors and diffusible crosslinkers (purple diamond heads). Motors and crosslinkers do not bind or unbind but may slide along microtubules. Motors create tension in the linkers that hinder their progression, while promoting the hopping of crosslinkers heads and microtubule sliding. A steady state is reached where motors and crosslinkers move on average at the same speed towards the plus end. By impairing the movements of the crosslinkers (red crossed arrows), occupancy decreases the sliding speed. **(c)** System C has diffusible motors composed of a motor head (green circle) with a diffusible head (orange diamond). Tension generated by active motor movement is released by hopping of diffusible heads, and microtubule sliding. **(d)** System D has diffusible motors as described in (c) and diffusible crosslinkers as described in (b). Diffusible heads of crosslinkers (purple diamonds) interfere with other molecules of the same category but not across category.

## 2 ASSUMPTIONS

The general assumptions are the same for all systems. Motor and crosslinking entities are made of two heads, binding to different microtubules. Unbound entities are uniformly distributed in space, and their heads can bind with equal rates *k*_*b*_ if they reach a microtubule. A bound head may unbind with constant rate *k*_*u*_. If one head is attached, the other head can attach to the other microtubule if it is overlapping at this position, also with rate *k*_*b*_. An entity bound to two microtubules exerts an elastic force of stiffness *κ* and zero resting length (Fig. 1a). At the time of second binding, the gap *δ* between the two heads is null, but if microtubules slide, a tension *f* = *κδ* will build up. This tension is relieved if the heads move appropriately along the microtubules, or if the microtubules slide relative to each other. The movement of the heads along the microtubule is affected by the tension *f* in the associated link, differently for motors and passive heads. We consider three types of heads (Table 1). **Motor heads** move continuously, since we are considering situations where jamming of motors does not occur. Attached motor heads move towards the plus-end with a speed *v*_*m*_ = *v*_0_ (1 − *f*_*m*_*/f*_*s*_), depending on the force against which the motor is pulling *f*_*m*_, its unloaded speed *v*_0_ and stall force *f*_*s*_. Thus, an antagonistic force reduces motor speed linearly, as shown experimentally [12]. We define *γ*_*m*_ = *f*_*s*_*/v*_0_, the characteristic drag coefficient of the motor head, such that *v*_*m*_ = *v*_0_ − *f*_*m*_*/γ*_*m*_. This equation determines the force-velocity relationship of the motor. We note that Kinesin-14 moves towards the minus end of microtubules, but as microtubule assembly dynamics are ignored here and only one type of motor is present, we can ignore microtubule polarity as the system is unchanged by swapping ‘plus’ and ‘minus’ throughout. **Non-diffusible passive heads** do not move along microtubules, and must unbind to relocate on a filament, releasing the associated linker tension immediately. We define *γ*_*c*_ = *κ/k*_*u*_, the effective drag coefficient of the crosslinkers. **Diffusible passive heads** are modelled following [9]. They bind at discrete sites on the microtubule lattice, separated by *a* = 8 nm. Passive heads can diffuse on this lattice by hopping to adjacent sites with a rate *k*_0_. However, a crosslinker head may not move to a position that is already occupied, nor step out of the microtubule at its ends. In the absence of external force, passive heads hop equally in both directions, undergoing pure 1D diffusion with a coefficient *D*_1_ = *k*_0_ *a*^2^. When the tension *f*_*d*_ in the linker between the heads builds up, the upstream (*k*^+^) and downstream (*k*^*-*^) rates differ. How these rates vary is not known, but thermodynamic consideration dictates that for any pair of states (a, b) with potential energies (*U*_*a*_, *U*_*b*_), the transition rates should satisfy Arrhenius law: *k*_*a*→*b*_*/k*_*b*→*a*_ = *e*^*ϵ*^ with *ϵ* = (*U*_*a*_− *U*_*b*_) */k*_*B*_*T*, and this is fulfilled by assuming [23]:

**TABLE I:**
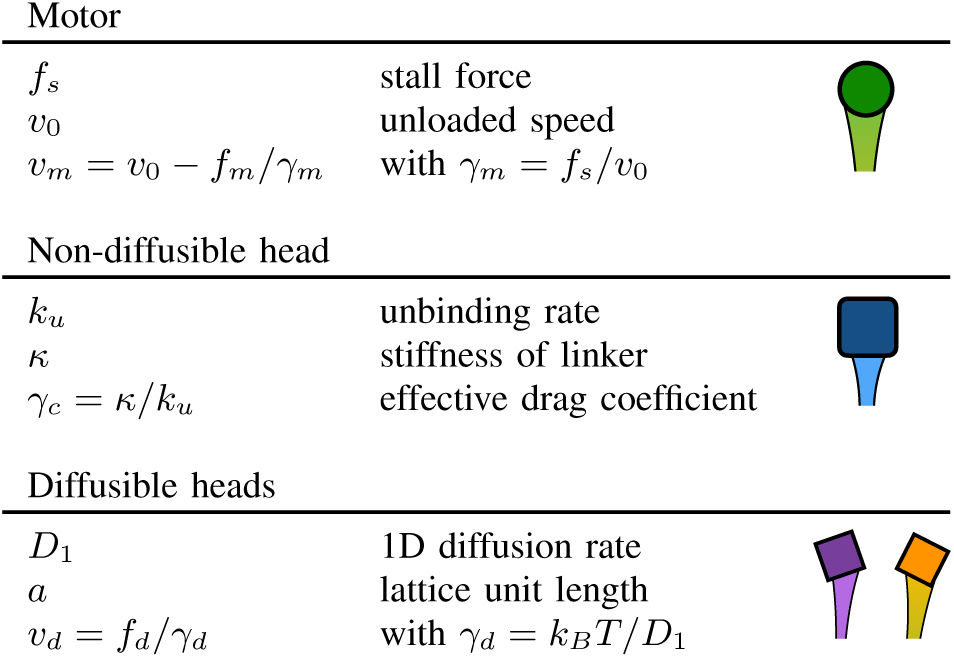
Microtubule binding heads.

| Motor |  |  |
| --- | --- | --- |
| $f_s$ | stall force | |
| $v_0$ | unloaded speed | |
| $v_m = v_0 - f_m/\gamma_m$ | with $\gamma_m = f_s/v_0$ | |
| Non-diffusible head |  |  |
| $k_u$ | unbinding rate | |
| $\kappa$ | stiffness of linker | |
| $\gamma_c = \kappa/k_u$ | effective drag coefficient | |
| Diffusible heads |  |  |
| $D_1$ | 1D diffusion rate | |
| $a$ | lattice unit length | |
| $v_d = f_d/\gamma_d$ | with $\gamma_d = k_B T/D_1$ | |

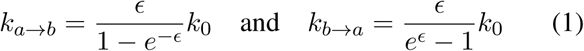

Since 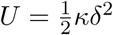, the hopping rates read:

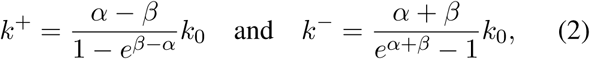

where *α* = *a f*_*d*_*/k*_*B*_*T* expresses the bias caused by force and *β* = *κa*^2^*/*2*k*_*B*_*T* echoes the difficulty of reaching a neighbouring binding site due to the stiffness of the linker. The diffusion rate of a crosslinker that is bound to two overlapping microtubules is defined by *a, k*_0_ and *β* and the microtubule’s own movements. In this article, we adopt the continuum limit that is obtained by neglecting the contribution of *β*. The drift speed along a microtubule, under a given force, then reads:

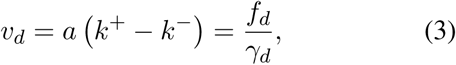

with *γ*_*d*_ = *k*_*B*_*T/k*_0_*a*^2^, the characteristic drag coefficient of a diffusible head. Accordingly, we adopted *κ* = 100 pN*/µ*m, a value for which the continuum limit is valid, for forces in the pN range (see discussion). **Microtubules** are incompressible lines oriented in opposite directions. The orientation of each microtubule dictates the natural movement of attached motors. The estimated viscous drag *γ*_fil_ ∼ 3*πξH/*[log(*H/d*) + 0.312] depends on the length of the microtubule *H*, its diameter *d* and the viscosity of the fluid *ξ*, following [22].

## 3 RESULTS

### 3.1 System A: Conventional motors and crosslinkers

We consider first non-diffusible crosslinkers that can bind and unbind from the antiparallel microtubules with constant rates *k*_*b*_ and *k*_*u*_ (Fig. 1a). The motors are of the Kinesin-5 family, sliding antiparallel microtubules apart. Given that all binding/unbinding rates and the overlap length are constant, there is a steady number of active motors in the system, producing an average force *F* between the microtubules. There is also a steady number of crosslinkers *c*, and their combined force must balance the motor force. The average force per crosslinker is then *f*_*d*_ = *F/c* = *κ δ*. Microtubules slide when one crosslinker unbinds, as the force of the motor is redistributed on a smaller number of crosslinkers. To evaluate the sliding associated with an unbinding event, we can consider the balance of forces after detachment: *F/*(*c -* 1) = *κ δ*^after^. The maximum displacement of the microtubule is therefore *δ*^after^ *δ* = *δ/*(*c* − 1). This will be the actual displacement, if the timescale of binding is sufficiently slow to allow the system to reach equilibrium, corresponding to *c k*_*b*_ *γ*_fil_ ≪ *κδ*. This condition holds true for realistic parameter values. In this regime, each filament will move at speed *v*_fil_ = *k*_*u*_*δ c/*(*c* −1), since 2*c* heads can unbind, and each unbinding event leads to a translation of both filaments by *δ/*2(*c* − 1). We can now calculate the force of the motors. Fast processive motors that are able to reach stall force (*v*_0_*κ/f*_*s*_ ≫*k*_*u*_, see table 2), will have reached a steady state defined by the force-velocity relationship *v*_*m*_*/v*_0_ = 1 − *f*_*m*_*/f*_*s*_. If *m* is the average number of bound motors, *f*_*m*_ = *F/m* = *κδc/m*, and by substituting *v*_fil_ we finally derive:

**TABLE II:** Parameters of simulations. Parameters used in the computer simulations, unless specified. † Binding and unbinding of bivalent motors is disabled in System B.

|  |  |  |  |
| --- | --- | --- | --- |
| Common | Linker stiffness<br>Lattice size | $\kappa$<br>$a$ | 100 pN/μm<br>8 nm |
| Rigid crosslinker | Unbinding rate<br>Binding rate | $k_u$<br>$k_b$ | 2.38 s<sup>-1</sup><br>1 s<sup>-1</sup> |
| Bivalent motor | Unbinding rate<br>Binding rate<br>Unloaded speed<br>Stall force | $k_u$<br>$k_b$<br>$v_0$<br>$f_s$ | $\dagger 0.01$ s<sup>-1</sup><br>$\dagger 0.01$ s<sup>-1</sup><br>0.05 μm/s<br>6 pN |
| Diffusive crosslinker | 1D Diffusion rate | $D_1$ | 0.1 μm<sup>2</sup>/s |
| Diffusive motor | Unloaded speed<br>Stall force<br>1D Diffusion rate | $v_0$<br>$f_s$<br>$D_1$ | 0.2 μm/s<br>6 pN<br>0.1 μm<sup>2</sup>/s |

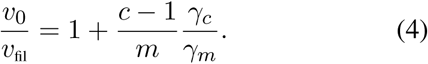

The filament speed (*v*_fil_*/v*_0_) is simply determined by the ratio of bound crosslinkers to motors (Fig. 2a) and the associated drag coefficients. As the system is symmetric, the filaments slide apart at speed 2*v*_fil_, not exceeding 2*v*_0_ as expected. Speed is independent of the absolute density of the molecules on the microtubule, and stable overlaps never form.

**Fig 2:**
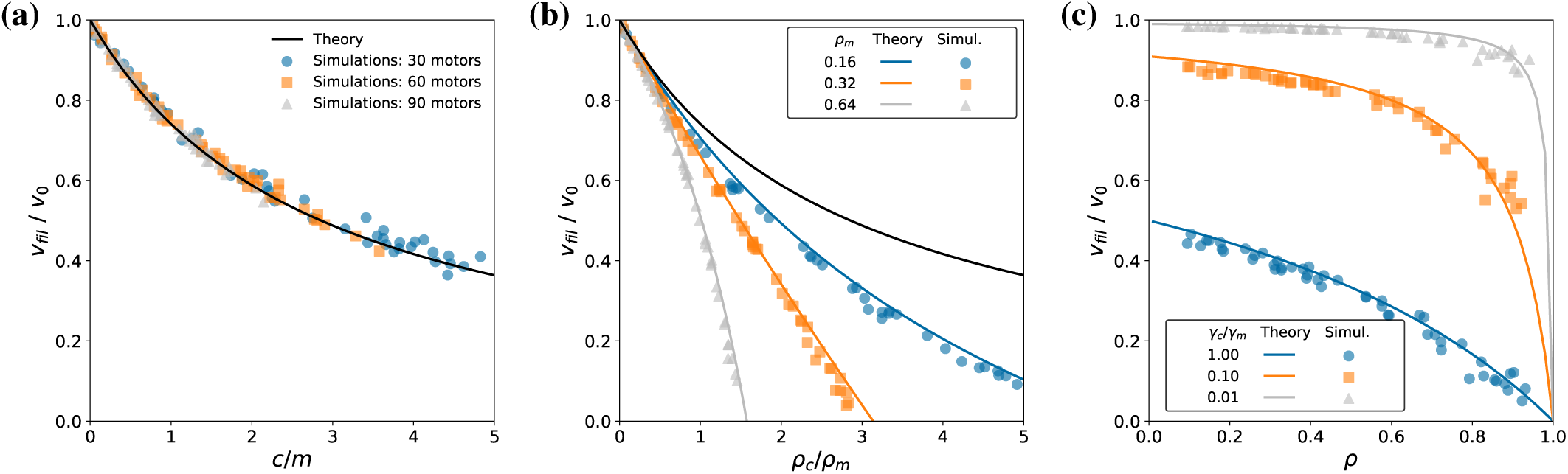
Systems A&B, bivalent motor and diffusible or non-diffusible crosslinkers. **(a)** Steady state speed for system A, with motors (*f*_*s*_ = 6 pN, *v*_0_ = 0.05 *µ*m*/s*) pulling non-diffusible crosslinkers (*κ* = 100 *pN/µ*m, *k*_*u*_ = 2.38 *s*^-1^), resulting in *γ*_*c*_*/γ*_*m*_ = 0.35. Dots represent the results of individual simulations containing 30 (blue circles), 60 (orange squares) or 90 (grey triangles) crosslinkers and a random number of motors (1 to 375). The line indicates Eq. 4. **(b)** Steady state speed for system B, with bivalent motors (*f*_*s*_ = 6 *pN*, *v*_0_ = 0.05 *µ*m*/s*) and diffusible crosslinkers (*D*_1_ = 10^-4^*µ*m^2^*/s*), resulting in *γ*_*d*_*/γ*_*m*_ = 0.35. Dots represent the results of individual simulations containing 60 (blue circles), 120 (orange squares), 240 (grey triangles) crosslinkers, and a random number of motors (1 to 380). Coloured lines show the corresponding predictions of Eq. 6. The black line represents the prediction for *ρ*_*c*_ = 0. **(c)** Steady state sliding speed for system B, varying *D*_1_ of crosslinkers. Dots represent the results of individual simulations, with *D*_1_ = 3.5 × 10^-5^ (blue discs), *D*_1_ = 3.5 × 10^*-4*^ (orange squares) and *D*_1_ = 3.5 ×10^-3^*µ*m^2^*/s* (grey triangles), resulting in *γ*_*d*_*/γ*_*m*_ = 1; 10^-1^and 10^-2^, respectively. These simulations included an equal amount of crosslinkers and motors, randomly chosen between 5 and 375. Since motors and crosslinkers do not unbind, the mean occupancies of crosslinkers and motors are equal. Coloured lines show the corresponding predictions of Eq. 6. *L* = 3*µ*m for all the simulations on this figure, and the horizontal and vertical positions of simulation dots are calculated from the simulation results (see methods).

### 3.2 System B: Bivalent motors and diffusible crosslinkers

We now consider diffusible crosslinkers, that however do not bind or unbind for simplicity (Fig. 1b). This setup is comparable to the PRC1/kinesin-5 system [20]. To account for the fact that only one crosslinker head may occupy each lattice site (*a* = 8 nm), we introduce the probability *ρ*_*c*_ ∈ [0, 1] for a lattice site to be occupied, and treat this value as if it was uniform along the lattice. In reality, since filament ends act as diffusion barriers, crosslinkers may accumulate at filament ends [1], [9]. However, in our case where the overlap is kept constant by microtubule growth, the crosslinkers remain equidistributed, and this parameter is effectively uniform.

Therefore, *ρ*_*c*_ = *c a/L*, and Eq. 3, becomes

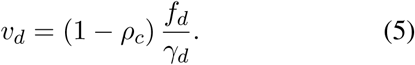

*In vitro*, the viscous drag of the filament is small compared to the drag of diffusible crosslinkers. For example, with *ξ* < 0.01 Pa.s, the viscous drag per unit length for microtubules is ∼ 0.015 pN.s.*µ*m^−2^ whereas *γ*_*d*_ = 0.04 pN.s.*µ*m^−1^ for *D*_1_ = 0.1 *µ*m^2^*/s*. Hence, at densities above 1 crosslinker/*µ*m, the force exerted by the viscous drag of the solution remains negligible, such that the force in the motor links should equal the force in the crosslinker links. With *m* motors and *c* crosslinkers, and calling *f*_*m*_ and *f*_*d*_ the forces per molecule, this means *F* = *m f*_*m*_ = *c f*_*d*_. In the steady state, because of the symmetry, motors and crosslinkers are immobile in space, and the speed of the heads is equal to the speed of the filament: *v*_fil_ = *v*_*m*_ = *v*_*d*_. Using the motor force-velocity relationships (*v*_*m*_ = *v*_0_ *-f*_*m*_*/γ*_*m*_) and Eq. 5, we derive:

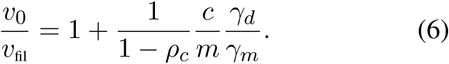

In addition to the ratio of motor to crosslinkers and their drag coefficients, the density of crosslinkers on the microtubule lattice also sets the filament speed (Fig. 2b). Higher occupancy leads to lower speeds (see Fig. 2b, black line obtained for *ρ*_*c*_ = 0). In the regime where the second term of the right hand side dominates, the speed is proportional to the number of motors. Sliding only stops when *ρ*_*c*_ = 1.

### 3.3 System C: Diffusible motors

We now consider diffusible motors composed of a motor head linked with a diffusible head (Table 1, Fig. 1c). We focus on the low density regime, and model the diffusible tail on a lattice without occupancy limits such that multiple heads can bind to the same site. A diffusible head unbinds immediately upon reaching the end of a filament.

At steady state, motors move towards the plus-end of their microtubule. Diffusible heads follow their motor at a distance *δ* behind, effectively moving towards the minus-end of the microtubule to which they are bound. With all links pulling in the same direction, the forces *f* of the links add up and the movement of the filament is *v*_fil_ = *m f/γ*_fil_, with *m* the number of links. The mean speeds of motor heads (*v*_*m*_) and diffusible heads (*v*_*d*_) are relative to their microtubules, which move in opposite directions, and the steady state requires *v*_*m*_ − *v*_fil_ = *v*_*d*_ +*v*_fil_. From the motor force-velocity and Eq. 3 we then derive *v*_0_ *-f/γ*_*m*_ = *f/γ*_*d*_ + 2*v*_fil_ and finally:

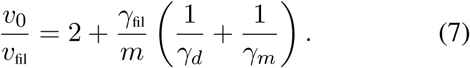

This formula can be compared to the sliding speed obtained in a gliding assay in which immobilised motors are pulling directly on the microtubule: *v*_0_*/v*_fil_ = 1 + *γ*_fil_*/*(*mγ*_*m*_). Firstly, the factor 2 in Eq. 7 indicates that diffusible motors, since they only contain one motor domain, can only slide microtubules at half their unloaded speed, unlike tetrameric kinesin-5 motors, which can slide microtubules at their unloaded speed. Secondly, part of the work produced by the motors is necessarily wasted in moving the diffusible head. Optimal microtubule transport is obtained for 1*/γ*_*d*_→ 0, but if the passive head can move, only a fraction of the motor force is transmitted to the link. This effect can be understood by considering immobile microtubules, for which the force in the link *f* is set by:

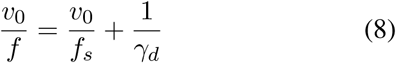

If *f*_*s*_ ≪ *v*_0_*γ*_*d*_ (the tail is hard to move), all the motor work is transmitted (Fig. 3a), but in any case the transmitted force is limited to *γ*_*d*_ *v*_0_ (Fig. 3a, dashed line). If *γ*_*d*_ *> γ*_*m*_, a significant fraction of the motor work will be used in sliding the diffusible head, rather than the microtubules. However, if *γ*_*d*_ ≪ *γ*_*m*_ (expected for the measured values), the force required to transport the diffusive tail on the microtubule is negligible compared to the stall force, and the motor heads move nearly at their unloaded speed. This means that the force produced is *γ*_*d*_ *v*_0_ (Fig. 3a, dashed line), corresponding to the drag force produced by diffusive tails moving at the motor’s unloaded speed.

**Fig 3:**
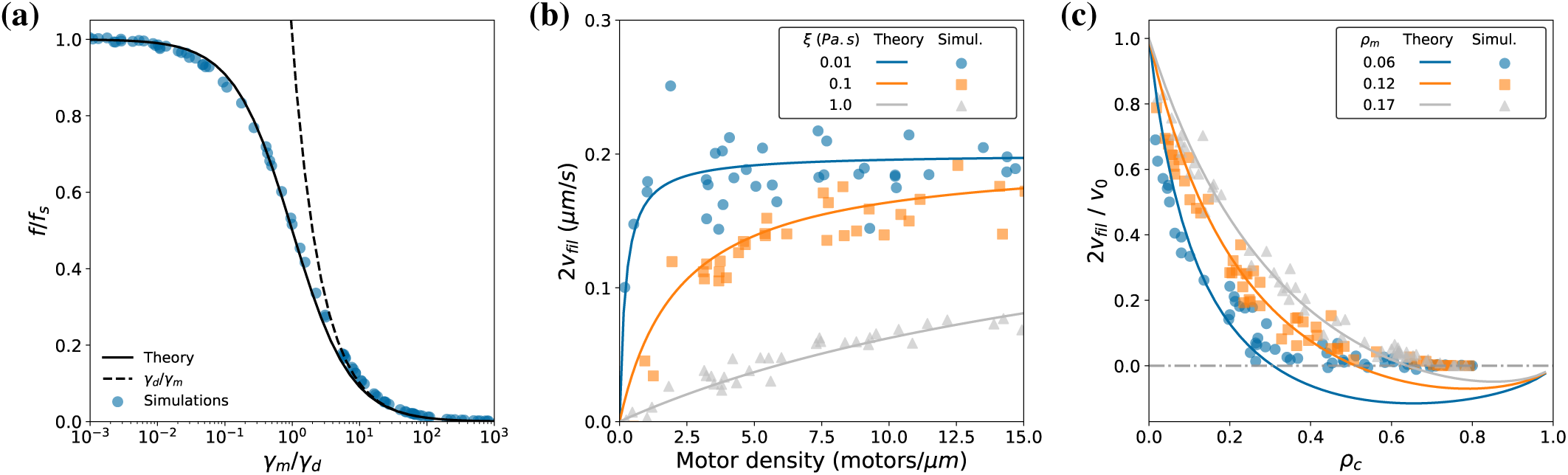
Systems C&D, sliding by diffusible motors. **(a)** The maximum usable force of a diffusible motor is limited by the drag coefficient of its diffusible head *γ*_*d*_. Dots represent the results of individual simulations with fixed microtubules and Kinesin-14 like motors (*f*_*s*_ = 6 *pN*, *v*_0_ = 0.2 *µ*m*/s*) and *D*_1_ ∈ [10^-7^, 1]*µ*m^2^*/s*. The line represents the prediction of Eq. 8. The dashed line represents the upper limit *γ*_*d*_*/γ*_*m*_. **(b)** Sliding speed for system C, with Ncd-like diffusible motors (*f*_*s*_ = 6 *pN*, *v*_0_ = 0.2 *µ*m*/s, D*_1_ = 0.1 *µ*m^2^*/s*), for different viscosities *ξ* in Pa.s: 0.01 (blue discs), 0.1 (orange squares) and 1 (grey triangles). Dots represent the results of individual simulations containing a random number of motors in [1, 100]. Coloured lines show the corresponding predictions of Eq. 7. **(c)** Sliding speed for system D, with diffusible motors, as in (b), and diffusible crosslinkers (*D*_1_ = 0.1 *µ*m^2^*/s*). Dots represent the results of individual simulations with varying number of motors: 100 (blue circles, *ρ*_*m*_ = 0.06), 200 (orange squares, *ρ*_*m*_ = 0.12), 300 (grey triangles, *ρ*_*m*_ = 0.18). The number of crosslinkers is randomly chosen in [1, 300]. Coloured lines show the corresponding predictions of Eq. 9. Note that simulations cannot yield negative speeds because overlap is kept constant by growth. For all simulations, *L* = 3*µ*m. All dots are placed according to the values of the relevant quantities averaged after the system has reached steady state (see methods).

System C was simulated for a motor with the characteristics of kinesin-14: *v*_0_ ∼ 0.2 *µ*m*/s, f*_*s*_ ∼ 5 pN [10] and *D*_1_ ∼ 0.1 *µ*m^2^*/s* [2], [14] (Fig. 3a, dots). We recover Eq. 7 and increasing viscosity reduces the sliding speed as anticipated (Fig. 3b). With *γ*_*d*_ = 0.04 pN.*s/µ*m and *γ*_*m*_ = 25 pN.*s/µ*m, certainly *γ*_*d*_ ≪ *γ*_*m*_ and the reduced sliding speed is set by *v*_0_*/v*_fil_ = 2 + *γ*_fil_*/*(*mγ*_*d*_). An interesting prediction can be derived from this formula. One can expect the drag of a filament to be roughly proportional to its length *H* (as predicted for *H >* 2 *µ*m [22]), and if the linear density of active motors is constant, the sliding speed will be independent of the length of the microtubule (since *γ*_fil_*/m* is constant). This is what has been experimentally observed [10].

### 3.4 System D: Diffusible motors and diffusible crosslinkers

We now add symmetric diffusible crosslinkers to system C (Fig. 1d). As shown experimentally, this slows down the sliding speed [1], [10]. In these experiments, one microtubule is fixed while an antiparallel shorter one is free to move and crosslinked by the diffusible crosslinker Ase1 and the motor Ncd. Sliding occurs at a constant speed set by the ratio of motors to crosslinkers. When the transported microtubule reaches the end of the fixed microtubule, the sliding stalls and eventually a stable overlap is established. During this time where the overlap decreases, the density of Ase1 increases but the density of Ncd remains unchanged. This suggests that Ncd turnover is faster than sliding, and that Ase1 does not compete with Ncd for binding sites. We make corresponding assumptions, with diffusible crosslinkers that do not unbind, and diffusible motors that bind and unbind with constant rates. Diffusible crosslinkers are modelled as in System B and the diffusible motors as in System C, and they do not interfere with each other for binding (Fig 1d). The diffusible heads from Ase1 and Ncd are distinct, and we note their drag coefficients *γ*_*d*_ and *γ*_*t*_ respectively (‘t’ for tail of Ncd). Given the observed parameters of kinesin-14 (*v*_0_ ∼ 0.2*µ*m*/s, D*_1_ ∼ 0.1*µ*m^2^*/s* [10], [2]) we expect forces produced by diffusible motors (Eq. 8, with *D*_1_ ∼ 0.1*µ*m^2^*/s*) to be in the same range as entropic pressure. The main force opposing the motor is thus the drag of the diffusible crosslinkers, as in System B, while the filament drag is negligible. We can use the contribution of the (positive) entropic pressure directly from [9]: *P* = −(*k*_*B*_*T/a*) log(1 *-ρ*_*c*_). The force balance becomes *c f*_*d*_ + *P* = *mf*_*m*_. We can calculate the filament sliding speed *v*_fil_ given that *v*_*d*_ = *v*_fil_ and *v*_*m*_ − *v*_fil_ = *v*_*t*_+*v*_fil_. Using Eq. 5, we obtain the following relation:

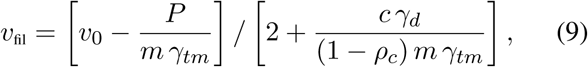

where we have defined 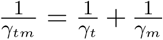. The denominator of the right hand side resembles the previous equations, while the numerator accounts for the entropic pressure. Speed decreases with crosslinker drag (*cγ*_*d*_) and increases with motor tail drag (*mγ*_*tm*_) as expected. For low densities (*ρ*_*c*_≪ 1), the sliding speed depends on the ratio of motors to crosslinkers [1]. Interestingly, the model predicts negative speeds if the entropic pressure is sufficient (Fig. 3c). Thus, stable overlaps may form for which *ρ*_*c*_ < 1. The result can be expressed from the density of species in the overlap *ρ*_*m*_ = *m a/L* and *ρ*_*c*_*= ca/L* as:

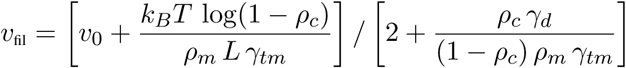

This reformulation highlights that the contribution of entropic pressure decreases with overlap length *L*, because it only depends on density, while the other forces exerted by motors and crosslinkers scale with *L*. Moreover, at high occupancies where *ρ*_*c*_∼ 1, the speed tends to zero as:

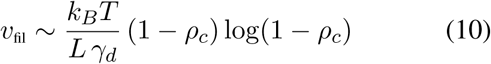

### 3.5 Entropic overlap expansion

We wondered if System D could recapitulate entropic overlap expansion, resulting from confinement of crosslinkers. This was measured experimentally by first applying hydrodynamic flow, to compress overlaps, and subsequently stopping the flow and measuring the expansion speed (Figure 3B from [9]). The expansion is purely driven by entropic forces, and can be analysed by omitting the motors from System D. We considered pairs of microtubules of length 20 *µ*m, with different initial overlap lengths. Assuming that the force under which these overlaps were formed is the same, the density of crosslinkers prior to the release of the force should be similar, since entropic forces depend only on density. Following these assumptions, the force per crosslinker is 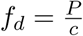 and using (2) we would predict a sliding speed:

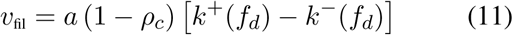

Assuming that *k*_0_ could be different on microtubule overlaps and on single microtubules, the measured diffusion constant on overlaps (0.011 *µm*^2^*/s*) can be matched by multiple combinations of *κ* and *k*_0_ (Fig. 4a). For such combinations, stochastic simulations show good agreement with the experimental data (Fig. 4b, c). For *κ <* 300pN*/µ*m the theory remains in good agreement with the stochastic model, provided that one uses Eqs. 2 to evaluate Eq. 11, taking into account the contribution of *β*. It seems that multiple parameter combinations could be adequate to model these results.

**Fig 4:**
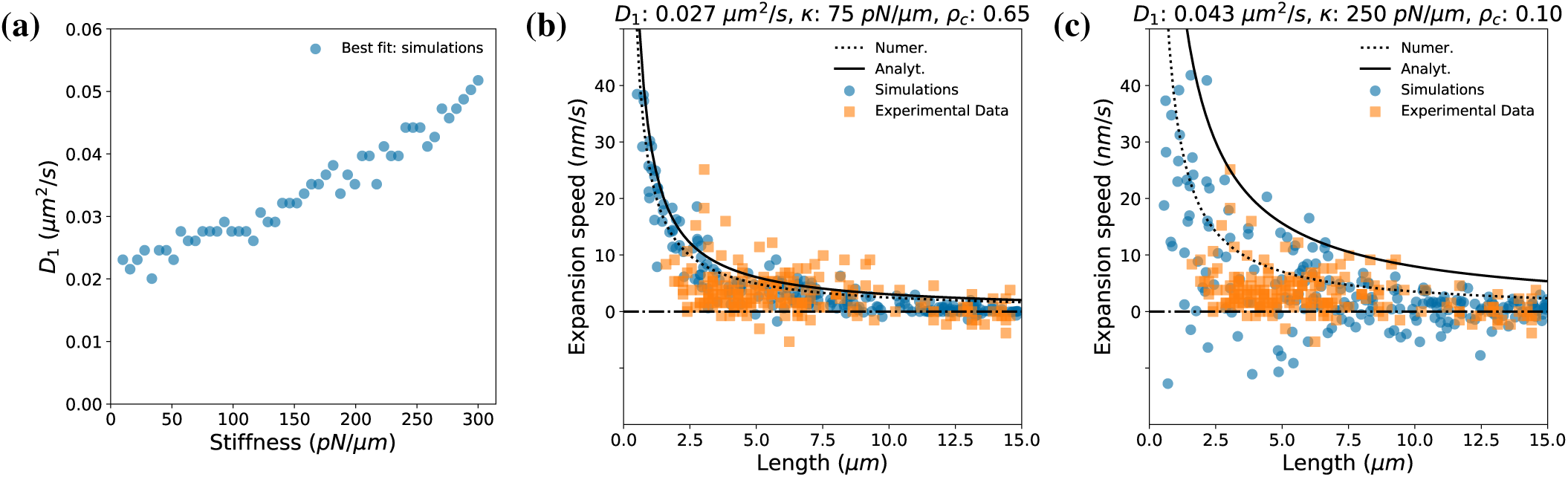
Entropic expansion of compressed overlaps. **(a)** Pairs of *D*_1_ and *κ* that best matched the observed diffusion rate of Ase1 in overlaps. For every value of *κ* (from 10 to 300 pN/*µ*m), simulations were run scanning 50 values for *D*_1_ (from 0.011 to 0.085 *µ*m^2^/s). A dot is placed indicating the *D*_1_ for which the simulated diffusion of 1000 crosslinkers was closest to the experimental value (0.011 *µ*m^2^/s). **(b)** Expansion speed for an overlap containing diffusible crosslinkers, as described in [9]. Blue dots represent the initial speed of sliding of individual simulations (*κ* = 75 pN*/µ*m), equilibrated with an occupancy *ρ*_*c*_ = 0.65. The black line represents the prediction of Eq. 11 with *β* ∼ 0, and the dotted line indicates the result obtained without this approximation. The experimental data (orange squares) is reproduced from [9] with permission. **(c)** Same as (b), but with a different value of the parameters: *κ* = 250 pN*/µ*m and *ρ*_*c*_ = 0.1.

### 3.6 Steady overlaps

In this last section, we consider the situation where sliding results in overlap shrinkage. Specifically, we aim to understand *in vitro* experiments that showed overlaps remaining for several minutes [1], [9], [26]. This phenomenon occurs when the turnover of crosslinkers is slower than sliding, such that crosslinkers accumulate in the overlap. On the contrary, if crosslinker turnover is sufficiently fast, the density of crosslinkers does not increase, and stable overlaps do not form [20]. Eq. 6 predicts that sliding stops when *ρ*_*c*_ ∼ 1. For values of *γ*_*m*_ ≪*γ*_*d*_ (the motors are stronger than the crosslinkers), the sliding indeed stops when crosslinkers are totally compacted at *L* = *c a* (Fig. 5a). However, for a diffusible motor, the entropic pressure can promote *L* > *c a* (Fig. 5b). From Eq. 9, an equilibrium between entropic pressure and motor force is reached if:

**Fig 5:**
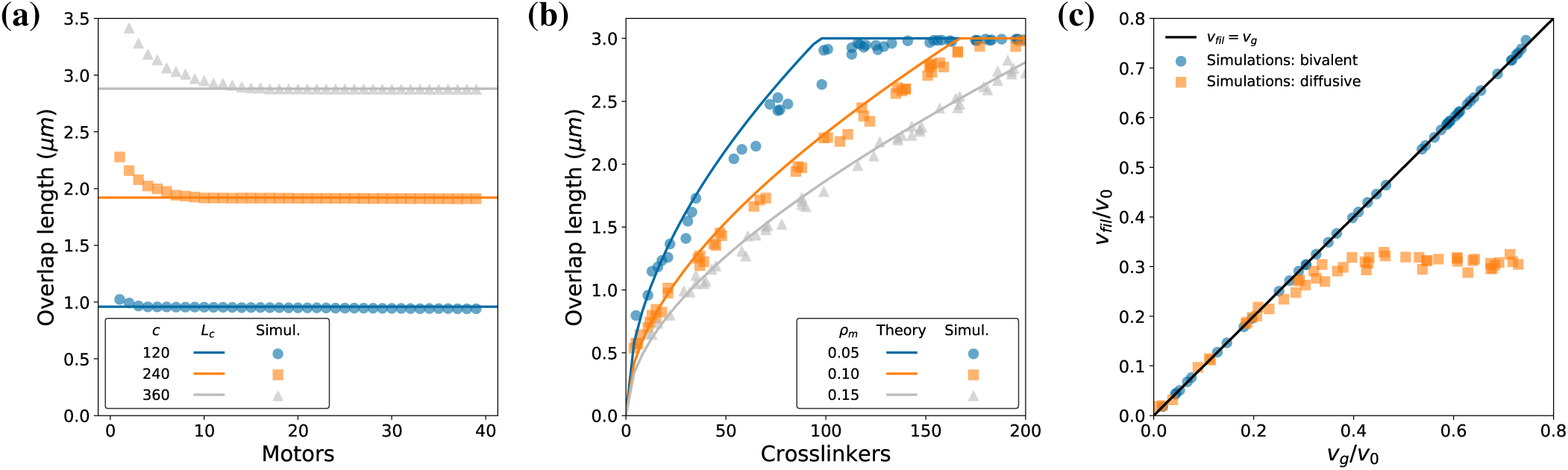
Steady state overlaps length and entropic forces. (a) Steady state overlap length for system B with non-growing microtubules. Dots represent the results of individual simulations, with 1 to 40 bivalent motors (*f*_*s*_ = 6 *pN*, *v*_0_ = 0.05 *µ*m*/s*). The number of crosslinkers (*D*_1_ = 0.1 *µ*m*/s*_2_) is 120 (blue discs), 240 (orange squares) and 360 (grey triangles). Coloured lines indicate total compaction. **(b)** Steady state overlap length for system D with non-growing microtubules. Dots represent the results of individual simulations, with 1 to 200 crosslinkers (*D*_1_ = 0.1*µ*m^2^*/s*). The number of motors (*D*_1_ = 0.1*µ*m^2^*/s, f*_*s*_ = 6*pN*) is 100 (blue discs), 200 (orange squares) and 300 (grey triangles) and *v*_0_ = 0.2*µ*m*/s*. Coloured lines indicate Eq. 12 and the dashed line indicates the predicted equilibrium obtained by not neglecting *β* in Eq. 2. **(c)** Sliding speed for systems B (blue circles) and D (orange squares), in which microtubules grow at a constant speed *v*_*g*_ (x-axis). Dots represent the results of individual simulations with 150 crosslinkers and 100 motors (blue) or 300 motors and 75 crosslinkers (orange). The parameters are as in (a) and (b). The black line indicates equality between growth and sliding speeds.

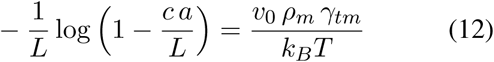

This result is confirmed by simulations (Fig. 5b), showing that even if entropic forces are smaller than the typical stall force of a single motor head, they are able to stabilise overlaps at densities above total compaction. Also, from Eqs. 6 and 9, we predict that, once a steady state length is reached, it can still decrease if crosslinkers unbind, or increase if more crosslinkers bind. Interestingly, diffusion rate of the crosslinkers does not affect final overlap length, but rather the speed at which this steady state is reached.

## 4 DISCUSSION

We have examined different ways by which motors and crosslinkers can be combined to make a stable overlap, predicting the sliding speed of the microtubules in each case. The analytical predictions matched the discrete stochastic simulations with *κ* = 100 pN*/µ*m. The equations resulting from the mean field approximation, without ignoring *β*, were solved numerically to improve the fit (Fig. 4c). However, for much higher values of *κ* and small forces per crosslinker, the system becomes qualitatively different. Microtubules adopt positions in which their lattices are in register, with an offset between them that is a multiple of the lattice unit. This regime was analyzed recently [25], showing how the jumping rate between two adjacent positions can depend exponentially on the number of crosslinkers. While the value of *κ* is critical in this model [25] as well as in ours, we note that the force may not be Hookean, such that measuring *κ* may be an ill-posed quest. In addition, the dependency of forward and backward rates on the force postulated in [9] is different from ours (Eq. 2), but both assumptions seem theoretically valid. Perhaps the most effective way to discriminate between these models is to directly determine the hopping rates of Ase1 under force.

The properties of the motors and diffusible crosslinkers operating in bundles are likely tuned for working together. Indeed, diffusible crosslinkers can regulate the sliding of diffusible motors, but they have little effects on the sliding caused by kinesin-5, even when they are in fair excess [20]. Given their biophysical characteristics, we can estimate if a motor would be hindered by crosslinkers or not. Kinesin-5 has a stall force of 1—10*pN* [5], and a speed of around 100 *nm/s* [5]. The diffusion rates of individual heads of kinesin-14 and Ase1 have been measured and they seem to be in the range of 0.1 − 0.01 *µ*m^2^*/s* [5]. From this, it appears that *γ*_*d*_ ≪*γ*_*m*_, suggesting that kinesin-5 motors would easily run over diffusible crosslinkers (Fig. 2c), which has indeed been observed [20]. Such strong motors can slide microtubules until the crosslinkers reach total compaction. Entropic pressure may be sufficient to stall less efficient force generators. Kinesin-14 (*v*_0_ ∼0.2 *µ*m*/s* and *f*_*s*_ ∼ 1*pN* [10]) has a diffusion rate that is comparable to Ase1/PRC1 diffusion. Thus *γ*_*m*_ ≪*γ*_*t*_, and we predict a significant effect on sliding speed, even at low occupancies (Fig. 3c). Thus many qualitative experimental observations are explained from the values of the parameters that have been published.

We have compared two types of molecular breaking: conventional crosslinkers that bind and unbind and diffusible heads. With the first type of breaking, sliding is determined by the ratio between motors and crosslinkers (Fig. 2a), while with the second type it depends also on the density of crosslinkers (Fig. 2b and 3c). Conventional crosslinkers do not sustain stable overlaps but diffusible crosslinkers can do so with both weak or strong motors. Motors like Kinesin-14 may stall against the entropic pressure (Fig. 5b) without compacting the crosslinkers completely. Motors such as Kinesin-5 would stop when crosslinker compaction prevents further sliding (Fig. 5a). System D (Fig. 4b, c) represents the experimental setups of [9], and reproduces qualitatively their main observations. Thus, while exponential friction [25] could explain the experimental expansion experiments [9], we propose here that considering lattice occupancy (Eq. 10) while adjusting parameters that are otherwise not constrained by experiments (Fig. 4) can also lead to the results observed. This alternative theory also explains that, under certain conditions, the sliding speed induced by kinesin-14 in the presence of Ase1 is independent of the overlap length, as observed in [1]. The same has been observed for kinesin-5 and PRC1 [20].

We have assumed that microtubules would grow at the required speed to maintain the overlap steady. We could however relax this assumption in simulations where microtubules were growing at a constant speed *v*_*g*_. They indeed reached a steady state overlap where growth and sliding equalise (Fig. 5c). A sharp reduction in speed at high densities of crosslinkers, as predicted for bivalent (Fig. 2c) and diffusible motors (Fig. 3c), is a key property for this synchronisation to happen. It allows for sliding to be conditioned on microtubule growth: microtubule elongation lowers the density of crosslinkers, and motors repack these crosslinkers by sliding the microtubules. Consequently, sliding and growing speeds will match without any further adjustment. The condition for this to spontaneously occur is that the motor/crosslinker system should be able to keep up with the required quantity of sliding: *v*_*g*_ should be slower than the maximum speed at which the microtubule can slide. Thus the appropriate equation (e.g. 6 or 9) are useful to estimate the conditions under which a stable overlap can be established.

We concluded that the timescale of crosslinker turnover was a critical parameter of the system. If turnover is faster than sliding, overlaps slide apart, as observed experimentally [20]. If turnover is slower than sliding, stable overlaps may form. Interestingly, fission yeast cells reduce the turnover of Ase1 upon anaphase entry, where maintenance of overlaps at the central spindle is important for the separation of spindle poles, suggesting that turnover regulation is active in cells [3]. Upon the action of only a few motors, the crosslinkers form stable overlaps, as observed experimentally [4]. The length of such overlap is determined by a simple rule: the overlap decreases until crosslinkers reach their maximal density. This is a remarkably simple and robust mechanism that does not require fine-tuned parameter values or intricate feedback loops. Thus, cells could adjust the length of the overlap by controlling the expression of crosslinkers.

Mechanically, with one molecular link every 8 nm over a few micrometers, such connections between two antiparallel microtubules are very strong. They are strong in the directions orthogonal to their main axis, as needed to maintain the two microtubules aligned. They are also strong in the axial direction, which is essential, particularly during anaphase where these overlaps contribute to spindle elongation. In contrast, any mechanism based on entropic pressure created by a ‘confined gaz of crosslinker’ is limited to relatively low forces and bound to result in high longitudinal compliance. This may or may not be desired depending on the operational demands placed upon the bundle.

In conclusion, perhaps the key property of these systems is to be able to accommodate microtubule assembly while maintaining steady and strong antiparallel connections, a conserved landmark of anaphase. While our theory is directly applicable to microtubules pairs formed *in vitro*, it will be valuable in the future to pursue geometrically realistic bundles made of more than two microtubules, to comprehend the mechanisms of action of diffusible crosslinkers *in vivo*.

## 5 ACKNOWLEDGMENTS

M. L-R is a fellow of DivIDE (https://divide-eunetwork.org). This project has received funding from the European Union’s Horizon 2020 research and innovation programme under the Marie Sklodowska-Curie grant agreement No 675737. F.J.N is supported by the Gatsby foundation. We thank S. Diez, M. Braun and T. Surrey for discussion and all the authors of [4] which inspired this work. We thank S. Dmitrieff for critical reading of the manuscript.

## 6 STOCHASTIC SIMULATION METHODS

We used the Open Source project Cytosim in 1D (www.github.com/nedelec/cytosim). The top (resp. bottom) microtubule is represented by an ordinate *p* (resp. *p*′) and a direction *d* = +1 (resp. *d*′ = −1). The location of the heads are recorded by their distance from the minus-end, a.k.a the abscissa *x*_*i*_, such that the position in space is *p*+*d x*_*i*_. An array of boolean values *T* is used for each microtubule to represent lattice occupancy, where *T* [*i*] corresponds to abscissas in [*ai, a*(*i*+ 1)]. The system is evolved using a time step of *τ* = 10^-5^*s*. Hopping to neighboring sites are stochastic events, simulated using a random number generator: a rate *R* is simulated by testing *θ <* 1 − *e*^*-Rτ*^ at every time step. Hopping is forbidden if the lattice is occupied, and the lattice is updated at each molecular binding, unbinding or displacement. The force in a link is *κ δ* with 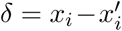 The movement of motors is represented by updating the abscissa: *x*_*i*_ = *x*_*i*_+*v τ*. The total force on each microtubule is calculated by summing all link forces. A Brownian dynamic approach using an overdamped Langevin equation is used to model the system, with an implicit numerical integration scheme [13]. **The steady state speeds** in Fig. 2, 3b, c were calculated from 40s of simulated time. The sliding speed was obtained by regression of the distance between the microtubule minus ends, from 8 to 40 seconds. Steady state speed measurements in Fig. 5c were calculated similarly from 100s of simulated time. The fitting for bivalent motors (blue dots) was done using data from 70 to 100s, and from 40 to 100s for diffusive motor (orange squares). **The steady state force** in Fig. 3a was measured from 40s of simulated time. The microtubules (as shown on Fig. 1c), were immobilized by a Hookean element of stiffness *κ*_*s*_. The steady state force is the average force exerted by these elements from 8 to 40 seconds. For each simulation, *κ*_*s*_ was adjusted to ensure that it would always have a similar stretch at steady state: *κ*_*s*_ ∞ *a*(*D*_*m*_*/v*_0_*kT* + 1*/f*_*s*_). **The steady state overlap length** for bivalent motors (Fig. 5a) was taken as the final overlap length after 100s of simulated time, while for diffusive motors, the average overlap length was calculated from 80 to 200 seconds. **The diffusion rate of crosslinker in overlaps** (Fig. 4a) was calculated from the mean squared displacement (*MSD/*2*t*) of 1000 crosslinkers bound to two microtubules after 1 second of simulated time, in simulations with an infinite capacity lattice. **The expansion speed** (Fig. 4b) was measured from 15 seconds of simulated time by regression of the distance between microtubule plus ends. All source code for simulation and analysis are available from the authors upon reasonable requests.

